# Tumor cell-organized fibronectin is required to maintain a dormant breast cancer population

**DOI:** 10.1101/686527

**Authors:** Lauren E. Barney, Christopher L. Hall, Alyssa D. Schwartz, Akia N. Parks, Christopher Sparages, Sualyneth Galarza, Manu O. Platt, Arthur M. Mercurio, Shelly R. Peyton

**Affiliations:** Department of Chemical Engineering, University of Massachusetts, Amherst, Amherst, MA 01003; The Wallace H. Coulter Department of Biomedical Engineering, Georgia Institute of Technology/Emory University, Atlanta, GA 30332; Department of Molecular, Cell and Cancer Biology, University of Massachusetts Medical School, Worcester, MA 01605

**Keywords:** Extracellular matrix, mechanotransduction, integrins, latency, biomaterials

## Abstract

Tumors can undergo long periods of dormancy, with cancer cells entering a largely quiescent, non-proliferative state before reactivation and outgrowth. For a patient, these post-remission tumors are often drug resistant and highly aggressive, resulting in poor prognosis. To understand the role of the extracellular matrix (ECM) in regulating tumor dormancy, we created an *in vitro* cell culture system that combines carefully controlled ECM substrates with nutrient deprivation to observe entrance *into* and exit *from* dormancy with live imaging. We saw that cell populations capable of surviving entrance into long-term dormancy were heterogeneous, containing quiescent, cell cycle arrested, and actively proliferating cells. Cell populations that endured extended periods of serum-deprivation-induced dormancy formed an organized, fibrillar fibronectin matrix via α_v_β_3_ and α_5_β_1_ integrin adhesion, ROCK-generated tension, and TGFβ2 stimulation. We surmised that the fibronectin matrix was primarily a mediator of cell survival, not proliferation, during the serum-deprivation stress, bacause cancer cell outgrowth after dormancy required MMP-2-mediated fibronectin degradation. Given the difficulty of animal models in observing entrance and exit from dormancy in real-time, we propose this approach as a new, *in vitro* method to study factors important in regulating dormancy, and we used it here to elucidate a role for fibronectin deposition and MMP activation.

## Introduction

Even after apparently successful treatment, disseminated tumor cells (DTCs) can remain viable for many years, entering into a prolonged period of dormancy before eventual reactivation and outgrowth. The presence of these disseminated, largely quiescent tumor cells in the bone marrow is a predictive marker of increased metastatic risk and poor prognosis [1, 2]. It has long been hypothesized that DTCs must land in hospitable tissues to grow, and many cells that disseminate to secondary organs are incapable, at least initially, of colonizing and proliferating into an overt metastasis [3]. However, it is debated whether the extracellular matrix (ECM) of the secondary tissue site, or any tumor-cell associated remodeling of said ECM, is a primary determinant of the fate of disseminated tumor cells.

Strong evidence is emerging to support that certain ECM proteins can promote cell dormancy, including thrombospondin 1 on the microvascular endothelium in breast cancer [4] and osteopontin within the bone marrow in leukemia [5]. In related examples, collagen I fibrosis is required for outgrowth of a dormant breast cancer population [6, 7], and collagen I-mediated signaling through DDR1 is required for reactivation of dormant breast cancer cells [8]. Physical properties of the ECM have been implicated as well, in that stiff environments promote cell proliferation, while softer surfaces support quiescence [9]. Related work has demonstrated that changes in the expression of ECM proteins can facilitate cell survival and outgrowth. For instance, fibronectin is produced by the stroma to support survival of metastasizing lung cancer cells [10]. Breast cancer cells have also been shown to trigger stromal periostin secretion [11] or produce their own tenascin C [12] to metastasize to the lung. Although not directly related to dormancy, these latter studies point to the importance of the generation of provisional matrices in first mediating survival, and then outgrowth of disseminated cells.

Although entrance into dormancy and reactivation are continuous and dynamic processes, most dormancy studies use static, endpoint measurements *in vivo*. Because of the difficulty of live imaging *in vivo* and the prohibitive cost of high-resolution sampling in time, these studies inherently cannot simultaneously capture the spatial dynamics and heterogeneity of microenvironment components alongside the ability to visualize the proliferation or quiescence of an entire cell population. An endpoint measurement such as immunohistochemistry can identify co-localized Ki67-labeled cells with ECM [4, 13], which correlates these proteins with proliferation. A major limitation with these studies is that there is no way to determine whether an individual Ki67-negative cell will eventually proliferate. Intravital imaging has started to address this problem, but it cannot yet view cells deep in the animal [5, 14-16], and it is nearly impossible to search an organ for an individual or small group of dormant cells. Despite these limitations, key recent *in vivo* studies have linked dormant cell outgrowth to fibrosis [6], TM4SF1 [8], miR-138 [17], and most recently laminin and smoking [14] in the lung. These efforts have required large pools of mice, particularly cost-prohibitive for most labs. We sought to find an alternative strategy that would enable higher throughput screens across cell sources and ECMs to determine how a population of disseminated cells enters dormancy and eventually transitions to overt metastatic outgrowth.

We developed a cell culture system capable of inducible and reversible population-level dormancy, where we can track the dynamics of individual cells and cell clusters. Using live imaging, we reveal significant heterogeneity across a dormant population of cells, simultaneously consisting of Ki67-positive, -negative, senescent, and actively proliferating cells. Across a diverse set of human breast cancer cell lines, those that could enter long periods of dormancy consistently produced and assembled a fibronectin matrix through α_5_β_1_ integrin-mediated adhesion and ROCK-mediated cell tension. The ability to proliferate after dormancy depended on the ability to degrade the fibronectin matrix with MMP-2. We demonstrated that the recipient ECM determined successful entry into and exit from dormancy, providing clues toward new ECM-associated therapeutic targets to prevent reactivation of dormant cell populations.

## Results

### In Vitro Model of Dormancy Reveals a Heterogeneous Population of Cell Cycle-Arrested and Proliferative Cells

We combined our established system of defined ECM protein surfaces [18, 19] with serum-deprivation to force cell populations into a dormant-like state (Fig. 1a). We define population level, or tumor dormancy (herein referred to as “dormancy” [20]) as a cell population that is not increasing in number, but remains viable for an extended time (at least 4 and up to 12+ weeks). The cell culture medium was serum- and growth factor-free, but contained amino acids, vitamins, sugars, and salts. Continuous culture under this serum-deprivation condition decreased both cell number and number of actively cycling cells (Ki67+) cells over the first 10-14 days, and, importantly, we could rescue cell growth (both total number and Ki67+) by re-introducing serum (ZR-75-1 cells shown as typical example in Fig. 1a). This is indicative of a reversible, quiescent phenotype in some subpopulation of cells within the dormant culture.

**Figure 1.**
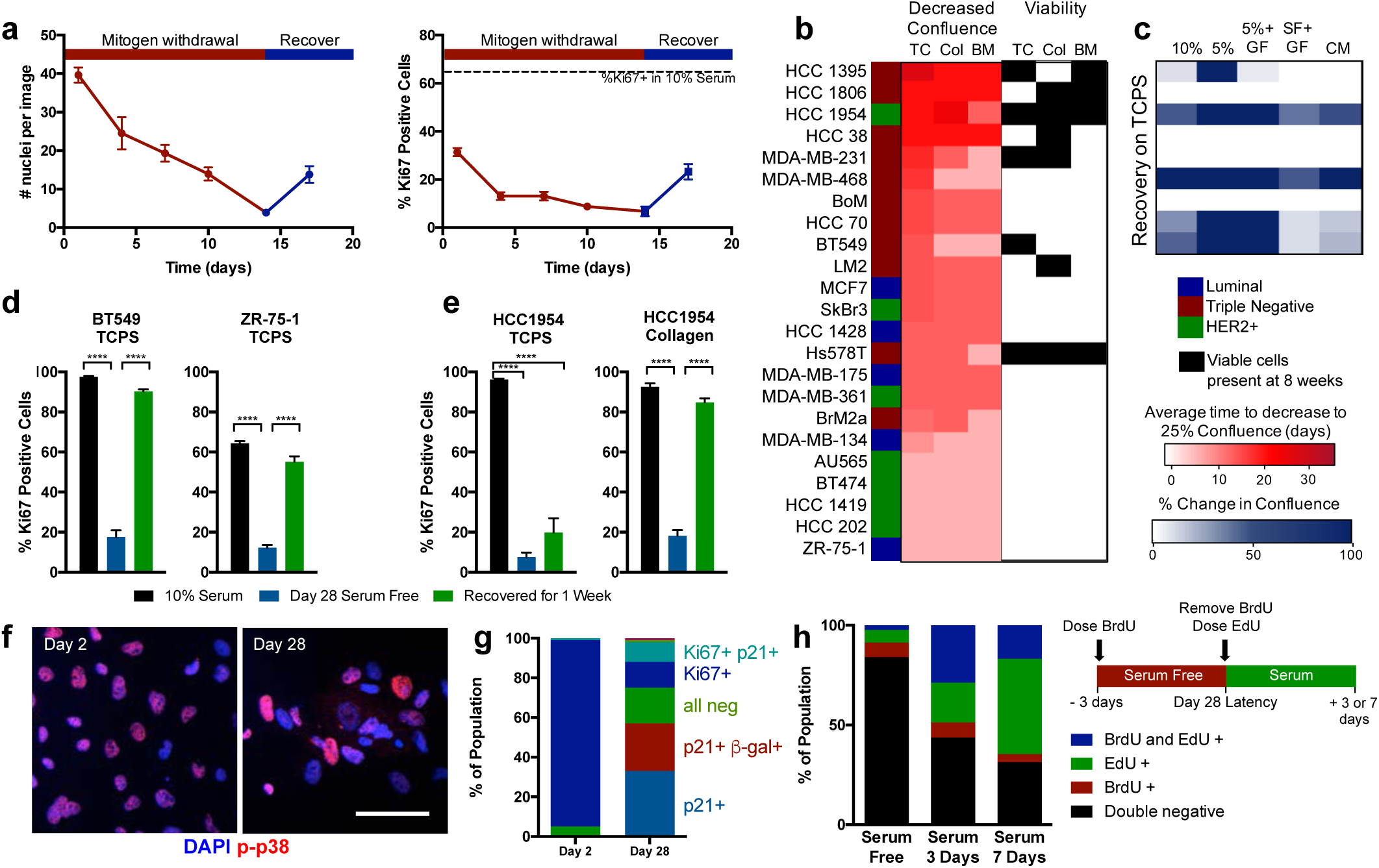
Serum withdrawal induces a reversible dormant phenotype in breast cancer cells. a) ZR-75-1 breast cancer cell line number and proliferation (Ki67 staining) over time with serum withdrawal (red) and recovery in full serum (blue). b) Survival of breast cell lines on TCPS (TC), collagen I coated coverslips (Col), or a mixture of ECM proteins inspired by those found in the bone marrow (BM)-coated coverslips. Average time to decrease confluence from 100% to 25% (scored by visual inspection by the same observer) is displayed on the left in red, and conditions where viable cells were detected at 8 weeks in culture are labeled on the right in black. c. Best performing cell lines were cultured on TCPS for 6 weeks, and stimulated with varying media conditions (GF: growth factor cocktail, CM: conditioned medium from MSCs). Heat map shows increase in confluence over 7 days stimulation in blue. d. Ki67 quantification of BT549, and ZR-75-1 plated on TCPS and e. of HCC1954 on TCPS or collagen. Black: 10% serum control, blue: day 28 serum-free culture, green: 7 day recovery. f. Immunofluorescent staining for phospho-p38 (red; Thr180/Tyr182) and DAPI (blue) in HCC1954 on collagen coverslips after 2 and 28 days of serum-starvation (Scale bar 100 μm). g. Population quantification of HCC 1954 via co-staining of Ki67, p21, and senescence-associated β-galactosidase. h. BrdU and EdU labeling experiment schematic and results for HCC 1954 on collagen. Black: double negative, red: BrdU-positive, green: EdU-positive, blue: double BrdU- and EdU-positive.. p < 0.05 was considered statistically significant. p < 0.05 is denoted with *, ≤ 0.01 with **, ≤ 0.001 with ***, and ≤ 0.0001 with ****.

We performed a screen on 23 human breast cancer cell lines to find those capable surviving 8 weeks of serum deprivation (Fig. 1b), and our ability to rescue growth after 7 days in serum or growth factor cocktails (Fig. 1c). Only 8 cell lines had viable cells remaining after 8 weeks of serum starvation, and of those, only 5 cell lines could be re-stimulated to grow. For some of the poor performing cell lines (e.g. ZR-75-1), we could boost their serum-deprivation survival by increasing the cell density (Fig. S1). We then asked if the ECM proteins presented on the cell culture surface affected this reversible dormancy by performing the assay on tissue culture plastic (TC), on a pure collagen 1 surface (Col), or on a mixture of proteins reflective of those found in the bone marrow (BM, a common site for clinical dormancy, Fig. 1b). Largely, the ability of the serum-deprivation stress to reduce cell culture confluency was independent of the ECM substrate, with few exceptions (most notably MDA-MB-468, BT549, MDA-MB-231). However, our ability to detect viable cells with a live-dead assay after a full 8 weeks of serum deprivation was highly substrate dependent for those cultures that did survive (Fig. 1b, right), with only the HCC1954 and Hs578T cell lines having viable cells left on all ECMs.

From this panel, we chose to further investigate the HCC1954, BT549, and ZR-75-1 cell lines, as these cell lines represent different clinical subtypes (HER2+, triple negative, and luminal, respectively), and they exhibited high, moderate, and low survival under serum starvation, respectively. Serum-free culture over 4 weeks induced population-level dormancy in these cell lines, defined by few Ki67+ cells (Fig. 1d-e). When stimulated with serum, each cell population recovered their proliferation within a week. Interestingly, the HCC1954 cell line only fully recovered the Ki67+ cell population on a collagen I-coated coverslip, not on TCPS (Fig. 1e). To more deeply examine whether ECM was important in promoting survival in this assay, we screened serum-free survival of all three cell lines across several ECM proteins and protein combinations, varying amounts of those proteins, and different substrate stiffnesses. Of all these variables, we found that ECM protein identity, specifically collagen I, most significantly promoted cell survival during serum-starvation (Fig. S2). For this reason, all subsequent experiments were performed using HCC1954 on this substrate unless otherwise noted.

After 4 weeks of serum withdrawal, 10-20% of cells were still Ki67+, so we asked whether cells were actively proliferating, cell cycle arrested, or more homogeneously quiescent (Fig. 1d-e). First, we observed that a subset of cells expressed active p38 after 28 days of serum deprivation (Fig. 1f), characteristic of a well-characterized dormancy program *in vivo* [21]. Second, time-lapse imaging identified a small population of proliferating cells undergoing mitosis (Movie S1). Third, we found that expression of p21 was significantly elevated in our dormant population, indicative of cell cycle arrest (Fig. S3c). When we co-stained the population of cells after 2 and 28 days of serum-deprivation for Ki67, p21, and senescence associated-β-galactosidase, we found most cells were Ki67+ after 2 days, but there was a richly diverse set of cell cycle states across the population after 28 days (Fig. 1g). Most of the population was simultaneously p21+ and Ki67-, indicating that a large fraction of the surviving population was both cell cycle arrested and non-proliferative. However, importantly, cells remained that were negative for all markers, or were Ki67+, and we speculated that some of these cells were responsible for maintaining cell numbers after long periods of serum-deprivation and possibly responsible for re-growth when serum was replenished in the “recovery” experiments.

To directly test whether individual cells could regain proliferation after dormancy, we sequentially labeled cells with BrdU at day 25 of serum-starvation, and EdU immediately upon serum-stimulated recovery, and followed them for either 3 or 7 additional days. We saw a large population of cells that not incorporate BrdU during serum deprivation, but did incorporate EdU during reactivation (Fig. 1h). Interestingly, when repeating this labeling experiment during sequential 72h periods of dormancy (i.e., label with BrdU during serum-starvation, then label with EdU during a second period of serum-starvation), we identified small populations (∼7%) that were positive for either BrdU or EdU, but not both (Fig. 1h, Serum 3 days), identifying small populations that were initially quiescent, then proliferative. Finally, the very small percentage of double positive cells decreased from day 3 to 7, reflective of the cells slowly cycling during serum deprivation, and able to regrow after serum reintroduction.

This method of serum deprivation-mediated dormancy allowed us to create a population of quiescent (Ki67-) cells, stable for 4-8 weeks (Fig. S3a). We wondered if these cells were a result of selection for cells that were robustly “dormancy-capable”. When we performed successive rounds of serum-removal and recovery on a single culture, we observed no change in the number of surviving or re-growing cells compared to a naïve population (Fig. S3b), suggesting that we recapitulated the heterogeneity of the original parental population upon re-growth.

### Fibronectin Secretion and Organization is Required for Extended Dormancy

We observed that many of the surviving cells had a spread, adherent phenotype, with some larger clusters of cells assembled into semi-adhered spheroids (Movie S1). This suggested to us that surviving dormant retained both strong cell-matrix and cell-cell adhesion. Since our initial screen identified ECM protein identity as a key factor in serum-free cell survival, we hypothesized that the dormant cell cultures may secrete ECM proteins to support survival. We performed LC-MS on HCC1954 samples enriched for ECM proteins at days 2 and 28 of serum-deprivation, and again after 1 week of recovery in full serum (Fig. S4a-b). We were excited to find many collagens, glycoproteins, and proteoglycans that were detectable at day 28, and subsequently disappeared when cell cultures were serum-recovered (Fig. S4a-b).

We were particularly interested in ECM proteins with known, strong roles in integrin-mediated adhesion, such as fibronectin, laminins, collagens, vitronectin, and osteopontin. Many of these were detected at day 28 of serum-free culture by LC-MS, and some of them were no longer detectable after recovery in serum (Figure S4b). When we looked more closely at these proteins via immunofluorescence, we saw that laminin (identified with a pan-laminin antibody), vitronectin, and osteopontin were all detected at increased levels during the dormant HCC 1954 culture, but all appeared intracellular by visual inspection (Fig. 2a). Serum starved cells can uptake ECM to promote survival [22], potentially explaining the intracellular distribution of these proteins we observed. Only fibronectin appeared to be deposited at high amounts at the day 28 time point (Fig. 2a), and we observed this fibronectin secretion phenotype consistently across all three cell lines we tested (Fig. S5a). In particular, the HCC1954 cells organized a dense fibronectin matrix over 28 days of dormancy, on both TCPS and collagen surfaces (Fig. S5b). They did not assemble fibronectin during 28 days of culture in serum-containing medium on any ECM, demonstrating that this secretion happens as a result of serum-starvation, not normal growth culture.

**Figure 2.**
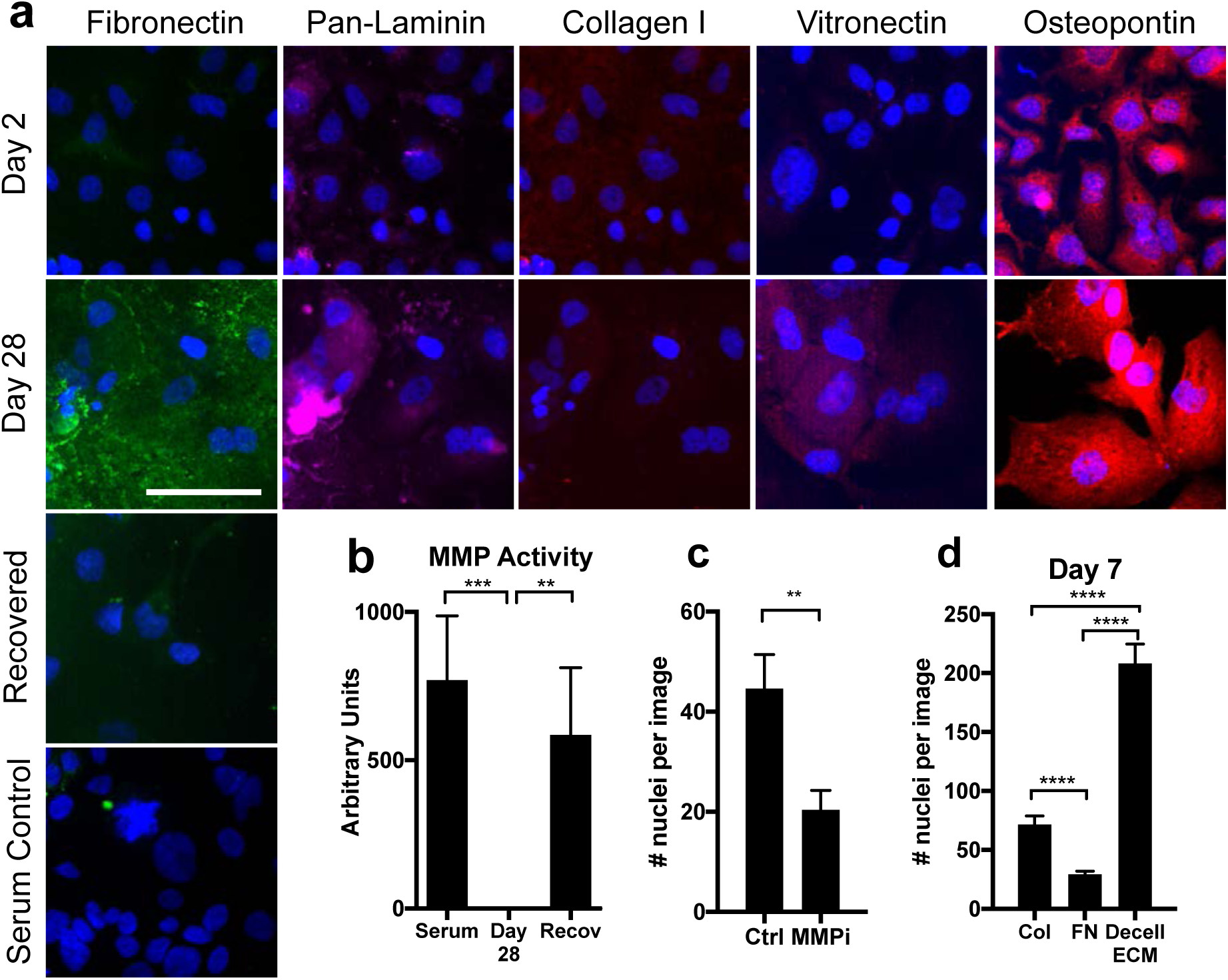
Dormant cells upregulate extracellular fibronectin specifically during serum withdrawal. a. Top rows, immunofluorescence for matrix proteins and DAPI in HCC 1954s on collagen in serum-free medium for 2 and 28 days. Bottom, immunofluorescence and for fibronectin in HCC 1954s grown for 28 days or serum starved for 28 days and recovered *in situ* for 1 week. Scale: 100 μm. b. Total MMP activity of HCC 1954 cells on collagen coverslips c. Number of nuclei resulting from reactivation of dormant cells treated with a pan MMP inhibitor for 7 days (MMPi; GM6001, 25μm) or control (serum-containing media, no inhibitor). d. HCC 1954 day 7 survival on collagen, fibronectin (FN), or HCC 1954 day 28 decellularized ECM (decell ECM) coverslips.

When the cells were cultured in serum-free conditions and then these same cells were recovered *in situ* with complete serum for 7 days, the fibronectin matrix was locally degraded where we saw high densities of cells (Fig. 2a, Fig. S6a). We hypothesized that cells were activating MMPs to degrade the fibronectin. Total MMP activity was measured with a fluorometric assay and MMP activity was high in cells under normal growth conditions, but undetectable in the dormant cell cultures (Fig. 2b). MMP activity was recovered with one week of recovery culture in serum, suggesting that MMPs were involved in the fibronectin degradation during reactivation. Specifically, zymography revealed that cathepsins S and V, and MMPs 2 and 9 were low, but detectable, in normal growth conditions. These decrease during the serum-free culture (with the exception of MMP-2), and cathepsin V and MMPs 2 and 9 were highly activated during growth recovery (Fig. S6b-c). Finally, when the cells were cultured the cells with a pan-MMP inhibitor (GM6001, 25 μM) during serum-stimulated recovery, HCC 1954 re-growth was reduced during serum recovery (Fig. 2c), suggesting that MMP activation and fibronectin degradation is beneficial for regrowth after dormancy. Interestingly, inhibition of cathepsin activity with E64 (5 μm) did not inhibit reactivation and outgrowth, suggesting that outgrowth is MMP-, but not cathepsin-dependent (Fig S6d).

To test whether the presence of a fibronectin-rich ECM directly supports long-term cell survival during serum-deprivation, we allowed the dormant HCC1954 cultures to assemble the insoluble fibronectin matrices over the 28 days of serum-deprivation, then we decellularized these samples, and reseeded new, naïve cell cultures onto them. When these cells were subjected to the long-term serum-deprivation on these decellularized, fibronectin-rich matrices, we found that HCC1954 and multiple other cell lines had significantly improved survival compared to the collagen surfaces across a several cell lines (Figs. 2d, S5c). This suggests that fibronectin promotes dormant cell survival during the serum starvation. However, some cell lines (AU565 and HCC1419) showed no ability to survive serum deprivation in the original screen, and also did not show improved survival with a decellularized ECM (Fig. S4c), so fibronectin is not sufficient to support entrance into dormancy by itself. We also surmised that the structure of fibronectin is important in this assay, because when we covalently coupled monomeric fibronectin to a coverslip, cells did not survive well (Fig. 2d). In fact, monomeric fibronectin did not support survival as well as collagen (Fig. 2d and S5d).

### Dormant Cells Assemble Fibronectin via Cancer-Cell Secreted TGFβ

Because matrix assembly is classically stimulated by TGFβ signaling, we explored whether TGFβ signaling affected the survival of dormant cells. When we provided cell cultures with exogenous TGFβ1 (1 ng/ml) or TGFβ2 (2 ng/ml) at every regular medium change, we saw no effect on cell survival at day 7 (Fig. 3a). Similarly, inhibition of TGFβ receptor signaling with LY-364947, a selective and ATP-competitive inhibitor of TGFβRI, did not affect day 7 survival (Fig. 3a). When the dormant cell cultures were exposed to the inhibitor over 28 days, however, cell survival was significantly decreased, and we observed a marked decrease in the fibronectin matrix (Fig. 3b-c). Continuous treatment of the cultures with exogenous TGFβ1 or TGFβ2 for the entire 28-day serum-deprivation culture either increased or maintained survival, respectively (Fig. 3b-c). Analysis of the culture supernatant demonstrated that HCC 1954 express undetectable levels of TGFβ1 and TGFβ3 (Fig. 3d), but high levels of TGFβ2 at day 28 of serum-free culture, suggesting that serum-starvation stimulates TGFβ2 secretion, which initiates fibronectin matrix remodeling and the survival of dormant cultures.

**Figure 3.**
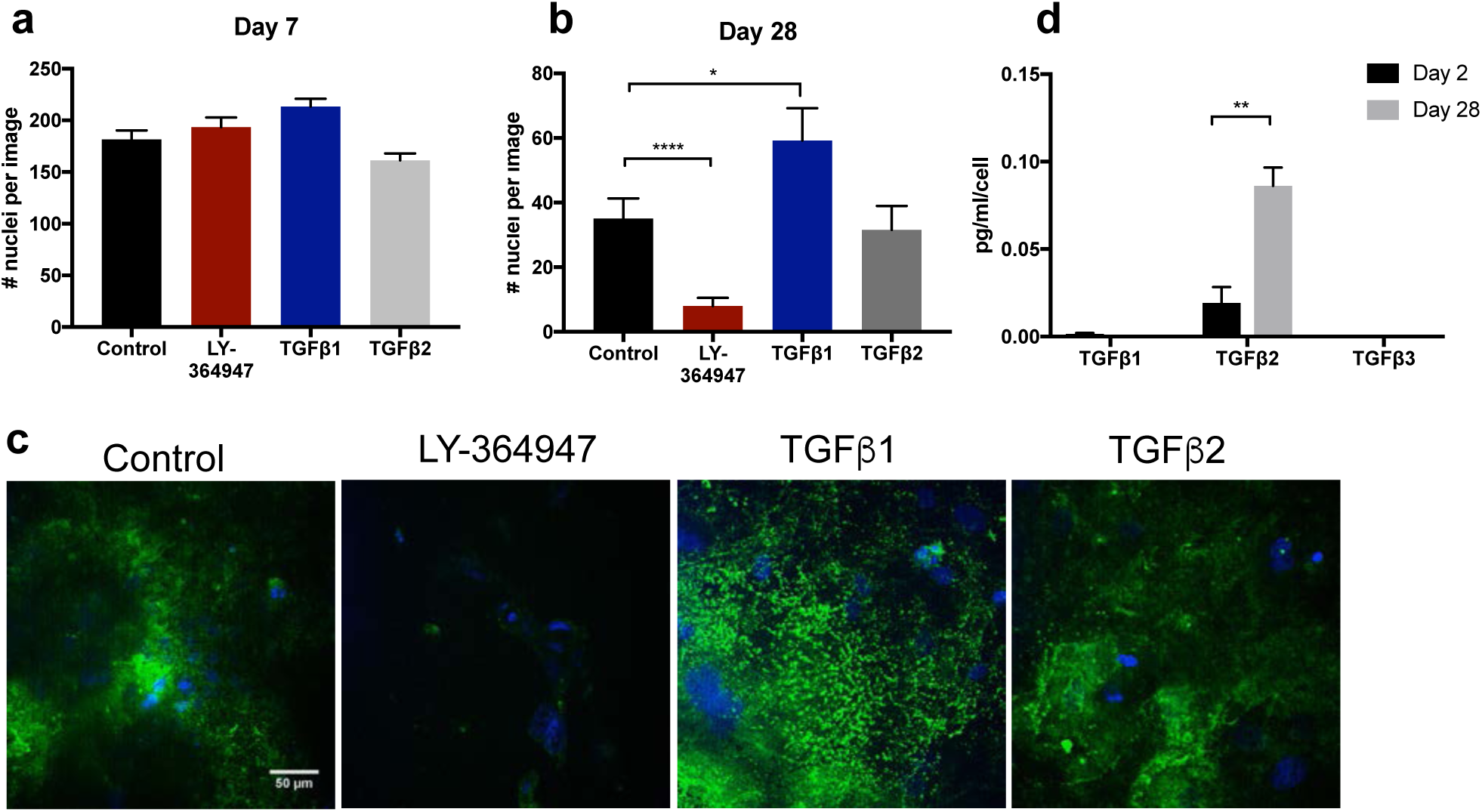
TGFβ stimulates fibronectin matrix production that mediates survival during mitogen withdrawal. a, b. Survival of HCC1954 under control, TGFβ receptor inhibition (LY-364947, 5 uM) and TGFβ1 (1 ng/ml) or TGFβ2 (2 ng/ml) stimulation over a. 7 days or b. 28 days. c. Immunofluorescence for fibronectin (green) and DAPI (blue) at day 28. d. Expression of secreted TGFβ1, TGFβ2 or TGFβ3 in serum-starved cultures at 2 and 28 days in culture. p < 0.05 is denoted with *, ≤ 0.01 with **, ≤ 0.001 with ***, and ≤ 0.0001 with ****.

### Fibronectin is assembled via α_5_β_1_ integrin-mediated tension, and mediates survival via adhesion through α_v_β_3_ and α_5_β_1_ integrins

Fibronectin assembly occurs via adhesion through α_5_β_1_ integrin and downstream RhoA activation, which then activates rho kinase (ROCK), generating tension to expose cryptic self-assembly sites in fibronectin, inducing polymerization [23-26]. The α_v_β_3_ and α_5_β_1_ integrin heterodimers bind to fibronectin with high affinity, and we observed these subunits were transcriptionally upregulated in our fibronectin-producing dormant cells (Fig. S7). We tested whether this pathway of fibronectin assembly was important for dormancy phenotype by inhibiting ROCK (Y-27632). We administered Y-27632 to cell cultures during serum deprivation during the first 7 days (entrance into dormancy), or continuously over all 28 days (long-term dormant survival), or, finally, we allowed the dormant cells to build up a fibronectin matrix for the first 21 days of dormancy, and added the inhibitors during days 21-28 (Fig. 4a).

**Figure 4.**
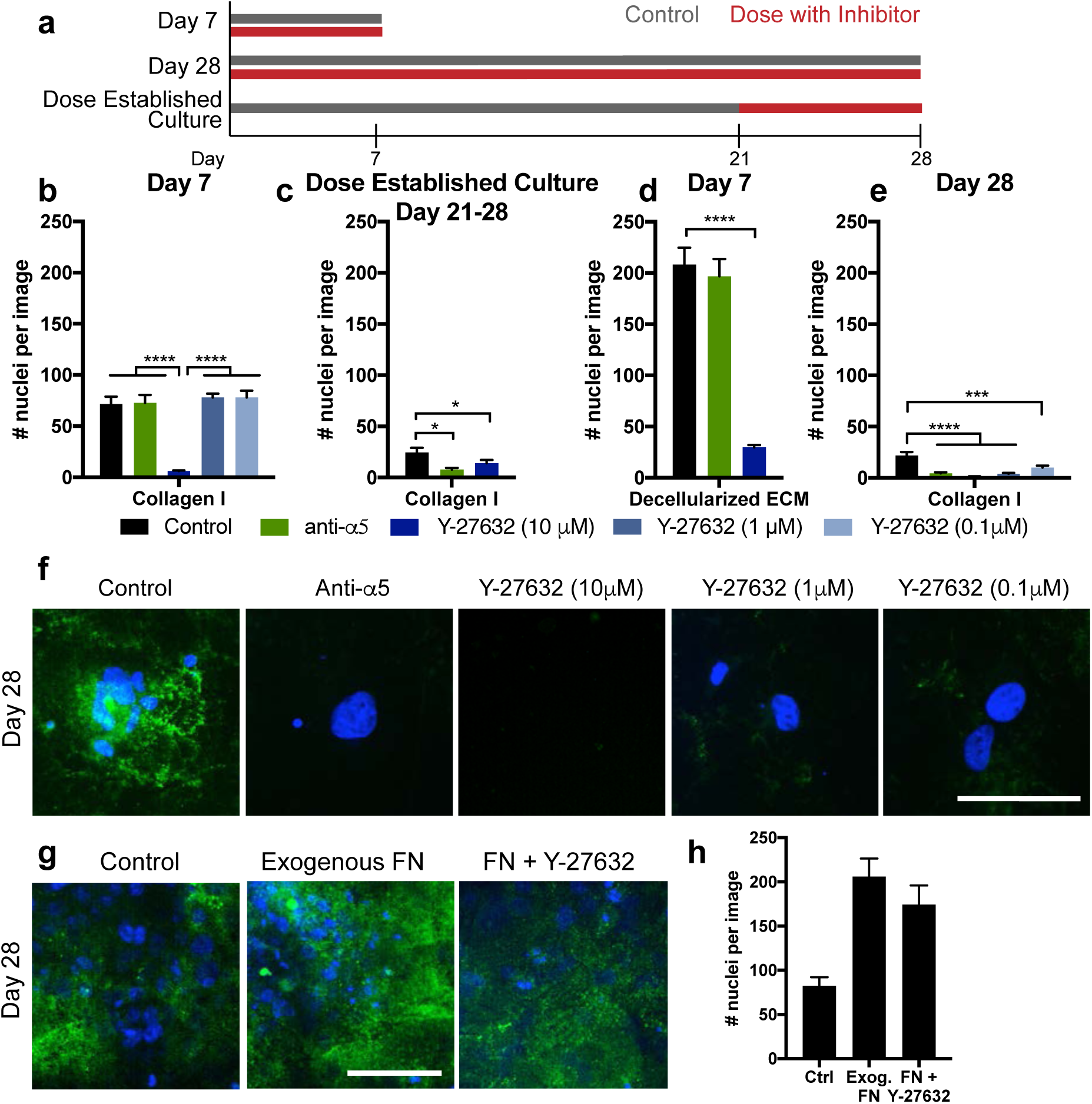
Dormancy-associated fibronectin is assembled via *α*_5_β_1_ integrin and ROCK to mediate survival. a. Experimental timeline of inhibitor dosing. Day 7 experiments were dosed continually through a day 7 endpoint; Day 28 experiments were dosed continually through a day 28 endpoint; separate cultures were established for 21 days, where inhibitor dosing was initiated for an additional 7 days (‘dose established culture’). b. Day 7 survival of HCC 1954 on collagen with inhibitors dosed at seeding and every medium change. c. Survival of cells with inhibitors, dosed after establishment of dormant culture for 21 days, and then subsequent dosing at every medium change through days 21-28. d. Day 7 survival of HCC 1954 on decellularized matrix with inhibitors dosed at every medium change. e. Day 28 survival of cells with inhibitors dosed at seeding and every medium change throughout the entirety of the experiment. Green: *α*_5_ integrin function affecting antibody; blues: Y-27632 (ROCK inhibitor at various concentrations). f. Immunofluorescence for fibronectin (green) and DAPI (blue) at day 28 on collagen with inhibitors dosed for the entire 28-day time period. Scale: 100 µm. g. Immunofluorescence for fibronectin (green) and DAPI (blue) at day 28 on collagen with exogenous fibronectin (10 µg/mL), or fibronectin with Y-27632 (1 µm). h. Day 28 survival of cells under control, exogenous fibronectin, or fibronectin with Y-27632. p < 0.05 is denoted with *, ≤ 0.01 with **, ≤ 0.001 with ***, and ≤ 0.0001 with ****.

The maximum dose of Y-27632 (10 μM) inhibited survival of cells over the first 7 days of serum-free culture on collagen-coated coverslips (Fig. 4b, dark blue bars). This maximum dose also inhibited cell survival when we administered the inhibitor after cells were allowed to pre-establish the fibronectin matrix (Fig. 4b-c), and when we seeded cells onto a decellularized ECM (Fig. 4d). However, 10- and 100-fold lower doses of Y-27632 did not affect survival during the first 7 days, but prevented survival of cells when administered over the full 28 days of serum deprivation (Fig. 4e). When we supplemented the culture with 10 μg/ml of soluble fibronectin to potentially jump-start the matrix, this promoted survival over the 28 days of serum-free culture, even in the presence of Y-27632 (Fig 4g-h). This suggests that cells use ROCK to secrete and assemble fibronectin to survive serum-deprivation culture. Maximum doses of ROCK inhibitor prevented cell survival in all cases, but these lower doses allowed us to observe the more specific role of ROCK in fibronectin assembly and long-term survival.

Inhibiting α_5_ integrin did not affect survival on collagen over the first 7 days (Fig. 4b, green bar), but it reduced cell survival when dosed after establishment of the fibronectin matrix (from day 21-28, Fig. 4c). When we seeded cells onto a decellularized matrix while inhibiting α_5_ integrin, we saw no change in survival over the first 7 days (Figure 4d). Finally, inhibiting α_5_ integrin function during the entire 28-day duration of the experiment inhibited cell survival under serum deprivation (Fig. 4e). We also saw an absence of fibronectin staining at day 28 in all these inhibitors conditions (Fig. 4f). Collectively, these results suggest that cells require α_5_ integrin to create the organized fibronectin matrices during serum deprivation.

When β_1_ integrin, which dimerizes with many alpha subunits (including α_5_), was similarly blocked there was minimal ability for cells to survive during serum-starvation, regardless of when we applied the treatment (including seeding cells onto decellularized matrices (Fig. 5b-d)). This demonstrates that β_1_ integrin was driving adhesion to both the collagen and fibronectin matrices. To more specifically target adhesion to fibronectin, we used cilengitide, a cyclic RGD drug that reduces fibronectin binding primarily via α_v_β_3_ and α_v_β_5_ integrins [27]. Cilengitide treatment did not affect the number of cells adhered to the collagen coverslip (Fig. 5b), but it was highly effective at disrupting survival when the dormant cells had assembled a fibronectin matrix (Fig. 5c), and when cells were seeded onto a decellularized matrices containing dense fibronectin (Fig. 5d). Although there is some evidence that α_v_β_3_ integrin can bind to denatured collagen [28], in our cells, cilengitide appeared to reduce adhesion to fibronectin but not to collagen-coupled coverslips.

**Figure 5.**
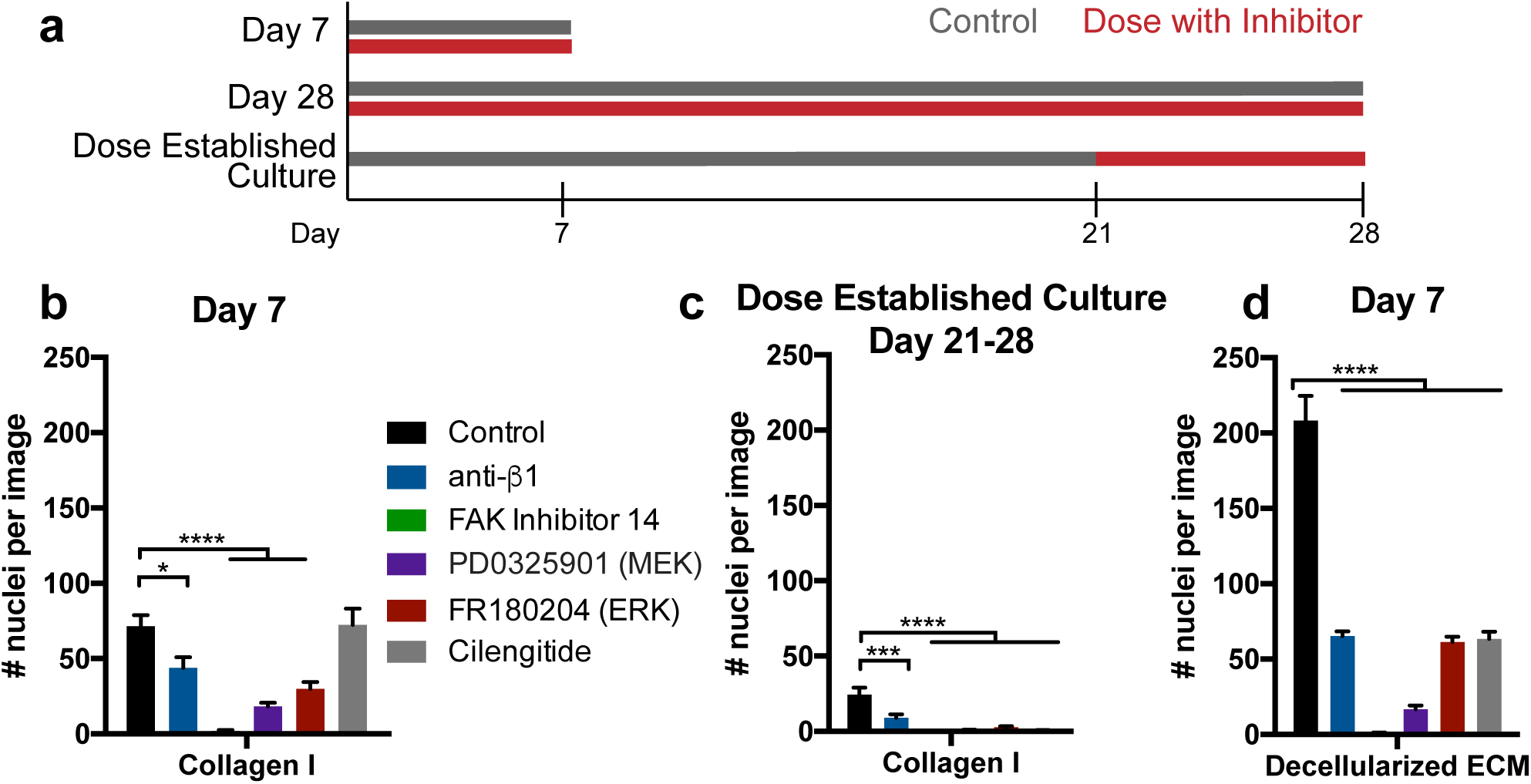
Survival is mediated via adhesion-FAK-ERK signaling. a. Experimental timeline of inhibitor dosing. Day 7 experiments were dosed continually through a day 7 endpoint; Day 28 experiments were dosed continually through a day 28 endpoint; separate cultures were established for 21 days, where inhibitor dosing was initiated for an additional 7 days (‘dose established culture’). b. HCC 1954 survival at day 7 on collagen coverslips with selected inhibitors dosed for the duration of the experiment. c. Survival of HCC 1954 cells after establishment of dormant culture for 21 days, and then subsequent dosing with inhibitors through days 21-28. d. HCC1954 survival at day 7 with inhibitors when seeded onto HCC 1954 decellularized ECM. Black: control, blue: anti-*β*_1_ (MAB17781, 1 µg/ml), green: FAK inhibitor 14 (10 µM), purple: PD0325901 (MEK inhibitor, 10 µM), red: FR180204 (ERK inhibitor 20µM), gray: cilengitide (100 µM). p < 0.05 is denoted with *, ≤ 0.01 with **, ≤ 0.001 with ***, and ≤ 0.0001 with ****.

We performed a small screen of tyrosine kinases to identify pathways activated during adhesion to the decellularized fibronectin matrices. Across this screen, only ERK was highly phosphorylated in both the dormant cell cultures and during adhesion to the decellularized ECMs (Fig. S8a-e). Pharmacological inhibitors to FAK, MEK and ERK all decreased cell survival over the first 7 days of serum starvation on both the collagen- and fibronectin-coupled coverslips and decellularized ECMs (Fig. 5b-d). This supports a hypothesis that cells are activating a survival pathway driven by α_v_β_3_ integrin mediated-adhesion to the assembled fibronectin matrix, downstream of ERK pathway activation during this serum-starvation induced dormancy in culture.

## Discussion

Applied cellular stresses, such as serum deprivation, application of drug, or hypoxia, have been used to select for cell populations primed for migration and metastasis [29-33]. By combining this approach with engineered materials, we selected for human breast cancer cell lines that can enter and survive dormancy based on their ability to secrete and organize ECM, a phenotype likely missed using traditional culture platforms or genetic profiling alone [34]. Our data supports the idea that dense fibronectin matrices facilitate long-term survival of cancer cells during serum deprivation-induced dormancy. Inhibiting α_5_β_1_, ROCK, or TGFβ receptor signaling prohibited fibronectin assembly, eliminating a large population of the surviving dormant cells. We suggest a role for α_5_β_1_ in this process, as adhesion to fibronectin through α_5_β_1_ integrin is associated with mature focal adhesions [23] to support intracellular tension, while adhesion through α_v_β_3_ does so weakly [35]. Reactivation and outgrowth of the cancer cells after the dormancy period could be inhibited by blocking their ability to degrade the fibronectin matrix via MMPs. Although they target different aspects of fibronectin matrix dynamics, both approaches could be promising mechanisms to eliminate dormant cell populations.

Other *in vitro* models have been used to study spontaneous dormancy [36, 37]; using far shorter culture times than our method here. Other approaches have used cell lines pre-selected with distinct proliferative or dormant potentials (e.g., the 4T1 and D2 series) [7, 17, 38-41]. Our study is unique in that a proliferative, heterogeneous cell population can be induced to *become* dormant via serum-deprivation stress, and the dormancy-competent cells can be subsequently reactivated. This approach is analogous to that used in the drug resistance studies, where resistant cells are selected via prolonged drug-induced stress [42, 43]. As one validation of our approach, the HCC 1954 cell line capable of showing this dormancy phenotype was also identified by Massagué and colleagues as one of two cell lines capable of generating dormancy-competent cells *in vivo* [13].

It has been proposed that dormant micrometastases are a mixture of arrested, quiescent, and slow cycling cells [44]. This has not been completely clarified since these studies were endpoint analyses, relying on defining dormant cells by fixed staining for lack of proliferation (Ki67-), DNA synthesis (EdU incorporation), or cell cycle stalling (p27+) [4, 21, 38]. These studies report that 70-80% of Ki67- or p27+ are dormant *in vivo* [4, 21]; however, in our culture system, the remaining 20-30% of Ki67+ cells could be cell cycle arrested (p21+) or proliferative (Figure 1g). By imaging live cells during the dormant culture, we revealed the presence of persisting cycling cells, quiescent cells, and arrested cells, a population-wide phenotype reminiscent of tumor dormancy.

We were initially surprised that our data appeared to contradict that of Aguirre-Ghiso, who has very thoroughly characterized an α_5_β_1_ integrin/EGFR/uPAR/fibronectin mechanism, mediating proliferation through ERK, not p38 [45, 46]. His lab showed that fibronectin fibrils block the p38 activity characteristic of quiescent cells, which results in sustained ERK signaling and proliferation [45, 47, 48]. Our data showed that dormant cells simultaneously suppressed p38 and activated ERK (Fig. S6a-b). However, when looking closer, immunofluorescence revealed that high phospho-ERK was ubiquitous (Fig. S6E), but phospho-p38 was heterogeneous (Fig. 1f). This suggests that this subpopulation of dormant cells may be activating the same stress signaling pathway described by the Aguirre-Ghiso lab [47]. In fibroblasts, serum withdrawal induces a reversible quiescence [49], which initially have decreased ERK activity that is eventually regained and sustained. This report is consistent with the high phospho-ERK we report at long dormancy times in cancer cells (Fig. S6a). Induction of stromal cell senescence is associated with acute activation of p38 and IL-6 release after genotoxic stress in lymphoma [50]. Therefore, it’s possible that the heterogeneous population of p38-high and –low cells is associated with senescent and quiescent cells, respectively, allowing for a population-level dormancy in response to serum-deprivation stress.

We found that the assembly of fibronectin coincided with entrance into a dormant phenotype, and this fibronectin assembly was dependent on TGFβ. Similarly, in cancer associated fibroblasts (CAFs), TGFβ is required to create a desmoplastic ECM [51]. We observed that cells required ROCK activation to assemble this fibronectin matrix, and other work has shown that there may be a feedback loop between tension-mediated TGFβ release and matrix assembly [52]. Cells responding to TGFβ are less proliferative, but more invasive and drug resistant [53]. Further, there are other reports of a TGFβ2 regulating the dormancy of tumor cells [21, 54]. The well-characterized senescence-associated secretory phenotype (SASP) is associated with a secretome that includes TGFβ [55]. This suggests that the senescent subpopulation we observed in our cultures (Fig. 1g) may be contributing to TGFβ release and subsequent fibronectin deposition. TGFβ is a promising target in many cancer clinical trials, and we suggest here a role for an anti-TGFβ therapy to prevent survival of dormant disseminated cells [56].

Our data suggests that targeting the TGFβ-mediated secretion of fibronectin allows cells to maintain a dormant state, agreeing with a recent demonstration that the optimal anti-TGFβ treatment for metastasis was during the pre-metastatic stage [57], although this is clinically challenging, because metastatic cells can disseminate extremely early during disease progression [58]. In our work, fibronectin disassembly occurred during the transition to outgrowth, and preventing MMP-mediated degradation of fibronectin may provide a better strategy to maintain dormancy and prevent reactivation. Together, this suggests that the dynamic control of fibronectin secretion/assembly and degradation may be a prerequisite for metastasis via promoting survival and subsequent outgrowth of dormant disseminated cells. Fibronectin expression, assembly, and degradation have all been featured as critical components of drug resistance and metastasis: MAPK and BRAF inhibitors can promote fibronectin production to promote drug resistance [59, 60]; serum starvation induces ECM secretion [61]; and hypoxia stimulates matrix remodeling and ensuing drug resistance [62]. Fibronectin, specifically, prevents serum deprivation-induced apoptosis via MEK1 and ERK activation [63], promotes proliferation [64], and permits survival of growth arrested cells through α_5_β_1_ integrin [65, 66], all in agreement with our findings.

We uncovered a mechanism by which dynamic fibronectin assembly and disassembly regulate breast cancer cell dormancy, which could be targeted to eliminate dormant cells or prevent their reactivation. Treatment regimens for eliminating dormant cells are difficult due to the stochastic nature of reactivation, and the extremely long time before relapse [67]; however, several treatment regimens have been proposed [67-70]. One camp argues that since these dormant cells are difficult to find and treat, they could be reactivated and sensitized to chemotherapy. This approach obviously comes with risk, as a patient would have awakened dormant cells that could be drug resistant. Another camp promotes a regimen that would keep cells in a dormant state, but this could be costly, lengthy, and unnecessary if a patient does not have micrometastases. A more exciting possibility is the recent revelation that dormant cells may in fact be sensitized to chemotherapy in an ECM-dependent manner [5, 71]. This agrees with our suggestion that targeting survival signaling from cell-ECM adhesion could be a valuable treatment option. There is promising preclinical evidence of efficacy of cilengitide as an anti-metastatic drug [72-74], and co-targeting dormant cells with cilengitide and chemotherapy may prevent metastatic outgrowth by eliminating dormant cells at the minimal residual disease stage.

## Materials and Methods

### Breast cancer cell culture

All cells were cultured at 37°C and with 5% CO_2_, unless otherwise noted. All cell culture supplies were purchased from Thermo Fisher Scientific (Waltham, MA). AU565, BT474, HCC 1395, HCC 1419, HCC 1428, HCC 1806, HCC 1954, HCC 202, HCC 38, HCC 70, ZR-75-1 were cultured in RPMI supplemented with 10% fetal bovine serum (FBS) and 1% penicillin/streptomycin (P/S). BT549, Hs578T, MCF7, MDA-MB-231, MDA-MB-468, MDA-MB-231 BoM (bone tropic), MDA-MB-231 BrM2a (brain tropic), MDA-MB-231 LM2 (lung tropic), SkBr3 were cultured in DMEM with 10% FBS, 1% P/S. MDA-MB-175 was cultured in Leibovitz’s L-15 medium with 10% FBS and 1% P/S without supplemental CO_2_, and MDA-MB-134 and MDA-MB-361 were cultured in Leibovitz’s L-15 medium with 20% FBS and 1% P/S without supplemental CO_2_. Tropic variants were a kind gift from Joan Massagué [75-77]; MDA-MB-231 were provided by Sallie Smith-Schneider; BT549, MCF7, and SkBr3 were provided by Shannon Hughes; and all others were provided by Mario Niepel.

### Biomaterial Preparation

Glass coverslips (5, 15, and 18mm) and PEG-PC hydrogels were prepared as previously described [18, 19]. Briefly, coverslips (Thermo Fisher Scientific) were UV-ozone treated and silanized through vapor phase deposition of (3-aminopropyl)triethoxysilane (Sigma-Aldrich) at 90 °C for a minimum of 18 hours. The coverslips were rinsed sequentially in toluene (Fisher Scientific), 95% ethanol (Pharmco-AAPER, Brookfield, CT), and water, and dried at 90 °C for one hour. They were then functionalized with 10 g/L N,N-disuccinimidyl carbonate (Sigma-Aldrich) and 5% v/v diisopropylethylamine (Sigma-Aldrich) in acetone (Thermo Fisher Scientific) at room temperature for two hours. Coverslips were rinsed three times in acetone and air-dried. ECM proteins (collagen I and natural mouse laminin were purchased from Life Technologies, human plasma fibronectin from Millipore, human collagen IV from Sigma-Aldrich, recombinant human tenascin C, osteopontin, thrombospondin-1, and vitronectin from R&D Systems, and laminin 511 and laminin 521 from Biolamina) were covalently bound to the glass surface by incubation at room temperature for three hours, then with 10 mg/cm^2^ MA(PEG)24 (Thermo Fisher Scientific) for two hours to block non-specific protein adsorption, rinsed three times in PBS, and UV-sterilized prior to cell seeding. Functionalized coverslips were placed into tissue culture plates that were coated with 100 μg/cm^2^ polyHEMA block non-specific cell adhesion.

For experiments on tissue culture polystyrene plates (TCPS), TCPS wells were coated with serum-containing medium at 37°C for at least 1 h prior to cell seeding to facilitate cell adhesion. ECM proteins were used at 2 Dg/cm^2^ unless otherwise noted. The Protein Atlas [78] was used to determine the composition of integrin-binding ECM proteins in human bone marrow, which primarily includes collagens, fibronectin, and laminins [79]. We created a minimalist representation of bone marrow ECM that contains proteins known to bind to cancer cells [18], composed of 50% collagen I, 15% laminin 521, 12% fibronectin, 8% tenascin C, 8% osteopontin, 5% laminin 511, and 2% vitronectin by weight. Collagen I and natural mouse laminin were purchased from Thermo Fisher Scientific (Carlsbad, CA), human plasma fibronectin from Millipore (Billerica, MA), human Collagen IV from Sigma-Aldrich (St. Louis, MO), recombinant human Tenascin C, osteopontin, thrombospondin-1, and vitronectin from R&D Systems (Minneapolis, MN), and Laminin 511 and Laminin 521 from Biolamina (Matawan, NJ).

### Establishment of Dormant Cell Cultures

Breast cancer cells were seeded onto biomaterials in serum-free medium using the specified culture medium for each cell line For seven-day experiments on different ECM proteins, protein concentrations, and hydrogels in Figs. 1a and S1, cells were seeded at 11,000 cells/cm^2^. For all other experiments, cells were seeded at 47,000 cells/cm^2^ to extend the possible duration. Serum-free medium was changed every 2-3 days for the duration of the experiment. At specified time points, cells were changed to serum-containing medium to grow a recovered population. Recovery was assessed after seven days of growth.

In cases where pharmacological inhibitors were used, inhibitor was replenished with every regular medium change. Concentrations of inhibitors and antibodies were chosen empirically to not affect initial seeding density. Where noted, inhibitors or antibodies were included in the serum-free or serum-containing medium at the following concentrations: E64, Tocris, 5 μm; ERK inhibitor FR180204, Sigma-Aldrich, 20 μm, FAK inhibitor 14, Sigma-Aldrich, 10 μm; soluble fibronectin, Millipore, 10 μ/ml; anti-α5 AF1864, R&D Systems, 1 μg/ml; cilengitide, Apex Biotechnology, 100 μM, anti-β1 MAB17781, R&D Systems, 1 μg/ml; MEK inhibitor PD0325901, Sigma-Aldrich, 10 μM; pan-MMP inhibitor GM6001, Millipore, 25 μM; ROCK inhibitor Y-27632, Millipore, 0.1, 1, 10 μM; pan-TGFβ AB-100-NA, R&D Systems, 20 μg/ml; anti-TGFBR1, LY-364947, Sigma-Aldrich, 5 μM).

Where noted in specific experiments, medium was supplemented with 1 ng/ml EGF (R&D Systems, Minneapolis, MN), 100 ng/ml FGF1 (R&D Systems), 100 ng/ml IGF1 (R&D Systems), 100 ng/ml HGF (R&D Systems), 1 ng/ml TGFβ1 (Sigma-Aldrich, St. Louis, MO), 100 ng/ml SDF1α (R&D Systems), or conditioned medium collected from mesenchymal stem cells to stimulate reactivation and growth. Where noted, medium was supplemented with a growth factor cocktail for recovery (100 ng/ml EGF (R&D Systems), 100 ng/ml FGF1 (R&D Systems), 100 ng/ml TGFβ1 (R&D Systems). In addition, 2 ng/ml TGFβ2 (Sigma-Aldrich) or 1 ng/ml TGFβ1 (Sigma-Aldrich) was supplemented in serum free medium.

### Characterization and Viability of Dormant Cell Cultures

Viability was assessed after 8 weeks of culture by staining with 4 μM ethidium homodimer-1 and 2 μM calcein AM (Thermo Fisher Scientific). Any condition where at least one viable cell was identified was considered positive for the presence of viable cells. Proliferation was assessed with immunofluorescence for Ki67, G1/S arrest by immunofluorescence for p21, and senescence by senescence-associated β-galactosidase (SA-β-gal) staining (Sigma-Aldrich). Immunofluorescence was performed according to standard protocols, and imaging was performed using a Zeiss Axio Observer Z1 with color and monochrome cameras or a Zeiss Spinning Disc Cell Observer SD (Zeiss, Oberkochen, Germany). The following primary antibodies were used for immunofluorescence: BrdU, ab8039, Abcam, 2 μg/ml; collagen I, ab6308, Abcam, 1:200; EdU, Click-iT EdU, Life Technologies, per manufacturer’s protocol; phosphor Trh202/Tyr204 ERK, #4370, Cell Signaling, 1:200; Fibronectin #563100, BD Biosciences, 1:200; Ki67, ab16667, Abcam, 1:200; pan-laminin, ab11575, Abcam, 1:100; osteopontin, ab8448, Abcam, 1:1000; phosphor Thr180-Tyr182 p38, #4511, Cell Signaling, 1:800; p21, ab7093, Abcam, 1:100; vitronectin, ab13413, Abcam, 1 μg/ml.

### BrdU and EdU Incorporation and Staining

10 μM BrdU (Thermo Fisher Scientific) was dosed for 3 days in serum free medium for 72 h prior to the day 28 endpoint. At day 28, the BrdU containing medium was removed, cells were washed once in serum free medium, and then stimulated with serum-containing or serum-free medium containing 10 μM EdU (Thermo Fisher Scientific). Cells were fixed after an additional 72 h or 7 days of culture, with regular medium changes. Cells were permeabilized in TBS-T, and DNA was denatured with 1M HCl (10 min, on ice), 2M HCL (10 min, RT), and phosphate-citric acid buffer (10 min, RT). Then, staining for EdU was performed according to the manufacturer’s protocol (Life Technologies), and, subsequently BrdU staining was performed using antibody labeling (Abcam, ab8039, 2 μg/ml).

### Decellularized ECMs

Decellularized ECMs were generated from dormant HCC 1954 cells cultures (originally seeded on collagen I-functionalized glass coverslips at 47,000 cells/cm^2^ in serum-free culture for 28 days) were according to a published protocol [80]. Briefly, cells were extracted with warm PBS containing 0.5% Triton X-100 (Thermo Fisher Scientific) and 20 mM NH_4_OH (Thermo Fisher Scientific), added drop-wise, and incubated at 37 °C for 10 minutes. Twice the volume PBS was then added, matrices stored at 4 °C overnight, and washed carefully with PBS 3 times. Decellularized ECMs were created and stored hydrated at 4 °C for up to 1 month prior to use. Decellularized ECMs were transferred to new well plates and rinsed once each in PBS and serum-free medium prior to use.

### Statistical Analysis

Statistical analysis was performed using Prism v6.0b. Data are reported as mean ± standard error. Statistical significance was evaluated using a one-way analysis of variance, followed by a Tukey’s post-test for pairwise comparisons. For Kaplan-Meier survival analysis, significance was determined using a Log-rank (Mantel-Cox) test. p < 0.05 is denoted with *, ≤ 0.01 with **, ≤ 0.001 with ***, and ≤ 0.0001 with ****; p ≥ 0.05 is considered not significant (‘ns’).

*Additional methods are described in the Supplemental Information, including antibodies for immunofluorescence and small molecule and antibody inhibitors or activators.*

## Author Contributions

LEB and SRP conceived the project and wrote the manuscript. LEB, MOP, AMM, and SRP designed research. LEB, CLH, ADS, ANP, CS and SG performed experiments. All authors discussed results and revised the manuscript.

## Acknowledgements

We thank Lauren Jansen, D. Joseph Jerry, and Mario Niepel for helpful discussions, and Lauren Jansen for providing a MATLAB code for quantification of cell number and the human bone marrow ECM analysis. We thank Joan Massagué, Shannon Hughes, Sallie Smith-Schneider, and Mario Niepel for generously providing cell lines, and Edna Cukierman and Sam Polio for assistance with the decellularization protocol. Mass spectral data were acquired at the University of Massachusetts Mass Spectrometry Core Facility, and we thank Dr. Stephen Eyles for assistance with sample preparation and data acquisition. This work was funded by an NSF-NCI award DMR-1234852 to SRP, and a grant from the NIH (1DP2CA186573-01). SRP is a Pew Biomedical Scholar supported by the Pew Charitable Trusts and was supported by a Barry and Afsaneh Siadat faculty award. LEB was partially supported by National Research Service Award T32 GM008515 from the National Institutes of Health. ADS was supported by a National Science Foundation Graduate Research Fellowship (1451512). ANP was supported by the Cell and Tissue Engineering NIH Biotechnology Training Grant (T32-GM008433), and MOP supported by a generous donation from the Giglio Family to the Wallace H. Coulter Department of Biomedical Engineering. AMM was supported by NCI R01 CA168464 and 203439.

