## Supplementary Materials for "Tumor cell-organized fibronectin is required to maintain a dormant breast cancer population"

**Supporting Information for Barney et al., “Cell-Organized Fibronectin is Required for Latency”**

**Supplemental Methods**

*PEG-PC Hydrogel Preparation*

PEG-PC hydrogels were prepared as previously described (1). Briefly, 17 wt% 2-methacryloyloxyethyl phosphorylcholine (Sigma-Aldrich) solution was prepared in PBS. Poly(ethylene glycol) dimethacrylate (Sigma-Aldrich) was added in concentrations ranging from 0.5% to 4% to create gels ranging from 1 to 64 kPa in modulus (1). The polymer solution was filtered through a 0.22 m filter, and bubbled with N_2_ gas for 30 seconds. A solution of 20 wt% Irgacure 2959 (BASF Ludwigshafen, Germany,) in 70% ethanol was added to the pre-polymer solution at 4 vol%. 70 l of this solution was sandwiched between a 3-(trimethoxysilyl)propyl-methacrylate treated coverslip and an untreated coverslip and exposed to UV light for 20 minutes. Gels were swelled for at least 48 hours in PBS, then treated with 0.3 mg/mL sulfo-SANPAH (ProteoChem, Hurricane, UT) in 50mM HEPES buffer at pH 8.5 and exposed to UV light for 10 minutes. Gels were then washed twice in PBS and flipped onto protein droplets containing 2 μg/cm^2^ collagen 1 for 18-24 hours. Gels were washed 2x in PBS and sterilized with germicidal UV for 1 hour before cell seeding.

*Proteomic analysis of ECM samples*

Cell-derived ECM proteins were obtained for serum free conditions day 2, and 28, and serum containing culture day 7. Samples were decellularized using 0.5% Triton in 20 mM ammonium hydroxide for 5 minutes, and rinsed two times with 1X PBS pH 7.4. Insoluble ECM proteins were enriched using the CNMCS compartmental protein extraction kit according to the manufacturer’s instructions (Millipore, Billerica, MA). This resulted in an insoluble ECM pellet.

The ECM-rich pellet left over from the CNMCS kit was solubilized and reduced in 8 M urea, 100 mM of ammonium bicarbonate, and 10 mM dithiothreitol (DTT) (Thermo) under agitation for 30 minutes at pH 8 and 37^0^C. After cooling, samples were alkylated with 25 mM iodoacetamide (Sigma-Aldrich) in the dark, and at room temperature for 30 minutes before the solution was quenched with 5 mM DTT. Subsequently, samples were diluted to 2 M urea with 100 mM ammonium bicarbonate at pH 8. Samples were digested with Lys-C endoproteinase (Promega, Madison, WI), at a ratio of 1:50 enzyme:protein for two hours under agitation at 37^0^C. Final digestion was performed in trypsin (Thermo) at a ratio of 1:50 enzyme:protein, under agitation overnight (12-16 hours) at 37^0^C, followed by a second aliquot of trypsin at a ratio of 1:100 enzyme:protein and two hours of incubation. Samples were desalted and concentrated using a C18 column (Thermo). A reverse phase Liquid Chromatography gradient was used to separate peptides prior to mass analysis. Mass spectrometry analysis was performed in an Orbitrap Fusion Tribrid (Thermo). Resulting peptides were aligned against the Matrisome using the Thermo Proteome Discoverer 1.41.14 (1). Parameters used trypsin as a protease, with 4 missed cleavages per peptide, a precursor mass tolerance of 10 ppm, and fragment tolerance of 0.6 Da.

*RT-PCR for Integrin and CDKN Expression*

Total RNA was isolated using the GenElute mammalian total RNA kit (Sigma-Aldrich) and 2.5 g total RNA was reverse transcribed using the RevertAid reverse transcription system (Thermo Fisher Scientific). 50 ng cDNA was then amplified using 10 pmol integrin-specific primers (Supplemental Table 3) and the Maxima Sybr green master mix (Thermo Fisher Scientific) on a Rotor-Gene Q thermocycler (Qiagen, Valencia, CA) as follows: 50 °C for 2 minutes, 95 °C for 10 minutes followed by 45 cycles at 95 °C for 10 seconds, 58 °C for 30 seconds, and 72 °C for 30 seconds. Both -actin and ribosomal protein S13 were included as reference genes to permit gene expression analysis using the 2^-ΔΔCt^ method. -actin, accession number: NM_001101.3 - 747F 5’ GGACTTCGAGCAAGAGATGG 3’; 980R 5’ AGCACTGTGTTGGCGTACAG 3’. RPS-13, accession number NM_001017.2 – 217F 5’ AAGTACGTTTTGTGACAGGCA 3’; 403R 5’ CGGTGAATCCGGCTCTCTATTAG 3’. Integrin and CDKN primers are provided in Supplemental Table 3.

*Phospho-Protein Quantification*

Coverslip samples for day 2 lysates and adhesion lysates were seeded at 47,000 cells/cm^2^, and for the adhesion lyates, both the adherent and suspension cells were lysed 15 minutes after seeding. Day 28 lysates were collected from 3-6 pooled samples seeded at 47,000 cells/cm^2^. Samples were washed once with ice cold PBS prior to lysis in RIPA buffer supplemented with protease (EDTA-free Protease Inhibitor Cocktail Tablets, 1 tablet in 10 ml, Roche, Indianapolis, IN) and phosphatase (1x phosphatase inhibitors cocktail-II, Boston Bioproducts, Boston, MA) inhibitors, 1 mM phenylmethylsulfonyl fluoride (Thermo Fisher Scientific), 5 μg/ml pepstatin A (Thermo Fisher Scientific), 10 μg/ml of leupeptin (Thermo Fisher Scientific), 1 mM sodium pyrophosphate (Thermo Fisher Scientific), 25 mM β-glycerophosphate (Santa Cruz Biotechnology, Dallas, TX). Protein concentration was determined with a BCA assay (Sigma-Aldrich). Lysate concentrations were adjusted to 400 μg/ml and were quantified with the MILLIPLEX MAP Multi-Pathway Magnetic Bead 9-Plex - Cell Signaling Multiplex Assay beads (Millipore) according to the manufacturer instructions, with the exception that beads and detection antibodies were diluted four-fold. Controls were performed to ensure that samples were within the linear dynamic range.

*Multiplex Cathepsin and MMP Zymography*

Medium was removed, and cells were washed once in ice cold PBS and then lysed in zymography lysis buffer (20 nM Tris-HCl at pH 7.5, 5 mM EGTA, 150 mM NaCl, 20 mM β-glycerol-phosphate, 10 mM NaF, 1 mM sodium orthovanadate, 1% Triton X-100, 0.1% Tween-20) with freshly added 0.1 mM leupeptin on ice. The multiplex gelatin zymography assay was performed as described previously to quantify the amount of active cathepsins and MMPs (2, 3). Briefly, 8 g of protein from each sample run in 12% (cathepsin) and 10% (MMP) SDS-polyacrylamide gels with an embedded gelatin substrate. Gels were then incubated in pH 6 cathepsin assay buffer (0.1 M sodium phosphate buffer, pH 6.0, 1 mM EDTA, and 2 mM DTT freshly added) or pH 7.4 MMP assay buffer (50 mM Tris-HCl at pH 7.4, 10 mM CaCl_2_, 50 mM NaCl and 0.05% Triton X-100) for 18 hrs at 37 °C. Gels were then stained with Coomassie Blue, destained to reveal cleared white bands, and then imaged with the ImageQuant LAS 4000 (GE Healthcare, Little Chalfont, United Kingdom).

**Supplemental Figures**

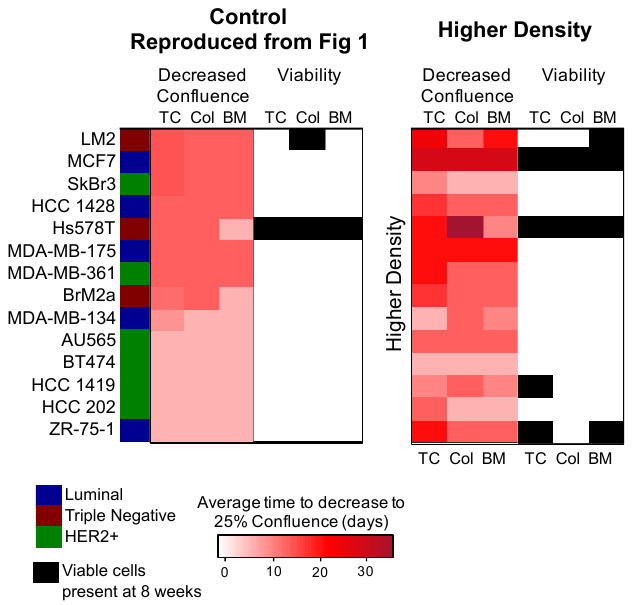

**Figure S1.** Heatmap of percent decrease in confluence and viability for bottom performing cell lines. Left, control density (data reproduced from Figure 1). Right, for higher cell density.

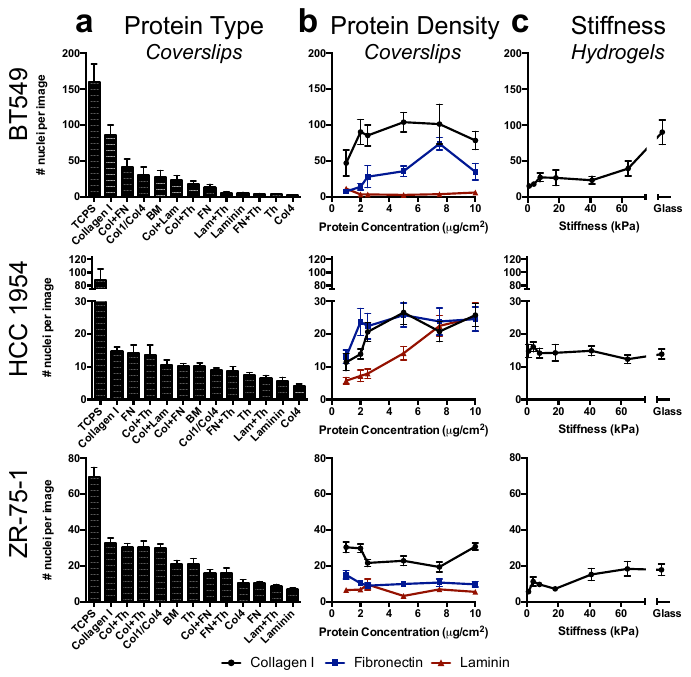

**Figure S2. Survival is maximized on collagen I coverslips.** Survival across varying matrix proteins at day 7 on coverslips (all at 2 μg/cm^2^). c. Survival across varying densities of collagen (black), fibronectin (blue), and laminin (red) at day 7 on coverslips. d. Survival across 2D PEG-PC hydrogels with varying stiffness (with 2 μg/cm^2^ collagen I).

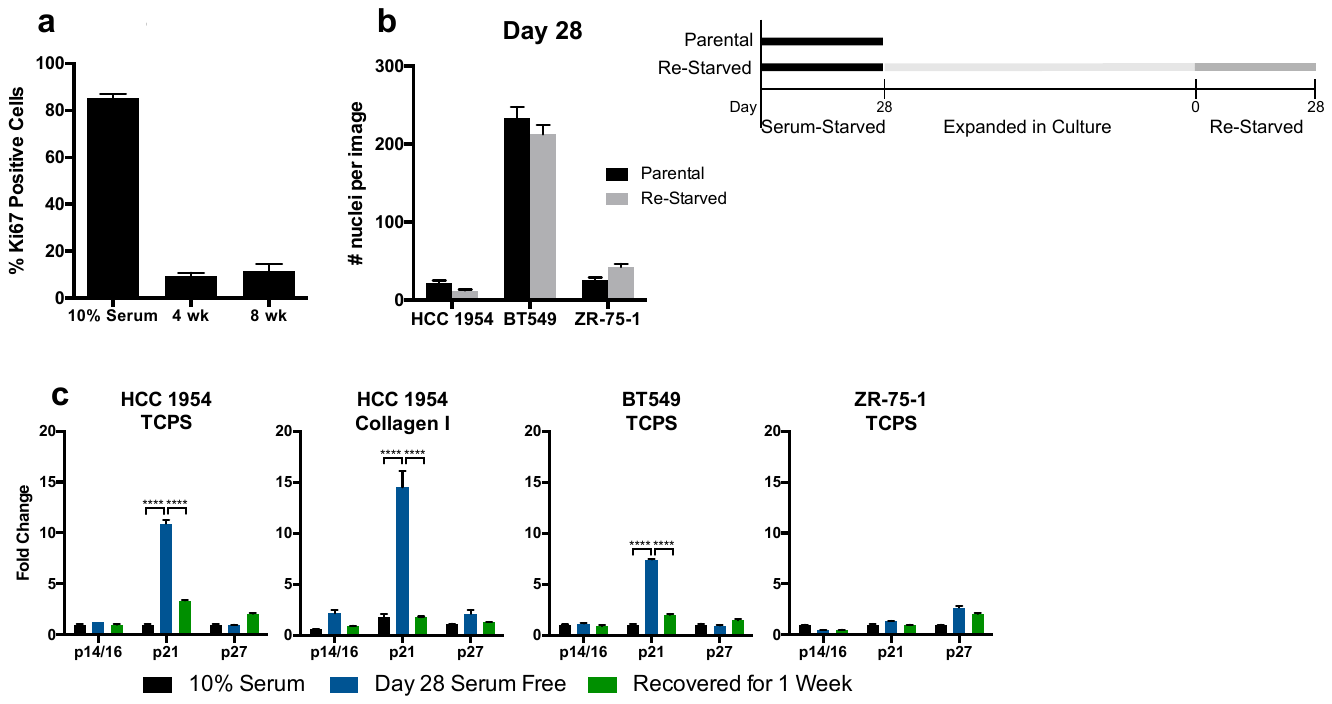

**Figure S3. Serum-starved cells are cell cycle arrested and non-proliferative.** a. Percent Ki67-positive HCC 1954 cells in serum, and after 4 and 8 weeks of mitogen withdrawal. b. Quantification of survival in parental and cell lines re-challenged with serum starvation a second time and timeline illustrating experimental procedure. c. RT-PCR for cell cycle inhibitors in HCC 1954, BT549, and ZR-75-1 on TCPS or collagen coverslips.

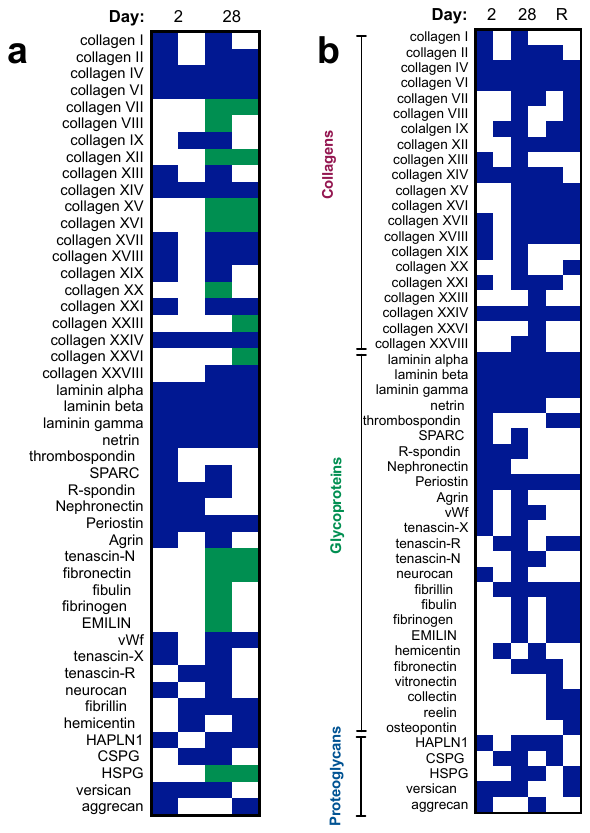

**Figure S4. Proteomic of extracellular matrix produced by HCC1954 during serum-starvation and recovery.** a. ECM produced by HCC1954 cells was produced and characterized by LC-MS at different time points and serum conditions. Core ECM proteins produced by the cells. Results were collected from two biological replicates. Proteins identified with at least 2 peptides were considered. Blue: peptide was detected, green: peptide was detected in day 28 but not day 2 samples. b. Comparison of peptides detected for day 2, day 28 and recovered samples.

**Figure S5. Dormant cells organize fibronectin.** a. Immunofluorescence for fibronectin in BT549 and ZR-75-1 on TCPS at day 2 or 28 in serum free culture. Scale: 100 μm. b. Immunofluorescence for fibronectin in HCC 1954 cells on TCPS or collagen during growth or 28 days of latency (serum starved). Scale: 100 μm. c. Survival of HCC 1954, ZR-75-1, MCF7, HCC 70, AU565, and HCC 1419 on collagen or HCC 1954 day 28 decellularized ECM. d. Survival of HCC 1954 cells on collagen and fibronectin coverslips at day 14 and day 28 of latency.

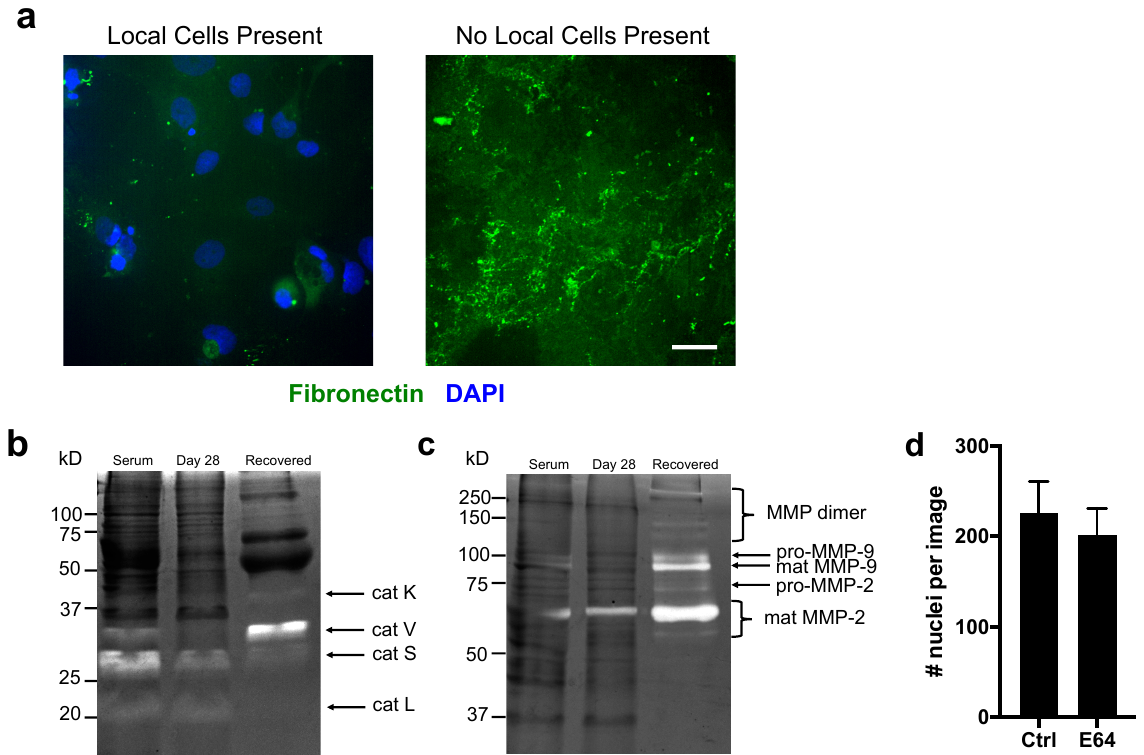

**Figure S6. Fibronectin is locally degraded during reactivation via MMPs**. a. Immunofluorescence for fibronectin (green) in latent HCC 1954 cells on collagen, recovered *in situ* for 7 days. Representative images are shown for areas on the same coverslip where recovered cells are present and absent. Scale: 100μm. b. Cathepsin zymography and c. MMP zymography for HCC 1954 cells on collagen. Serum: HCC 1954 in serum for 48 hours, day 28: dormant culture for 28 days, recovered: dormant cells recovered for 7 days. d. Recovery of cells serum-starved for 28 days and then recovered under control or cathepsin inhibition (E64, 5 μm).

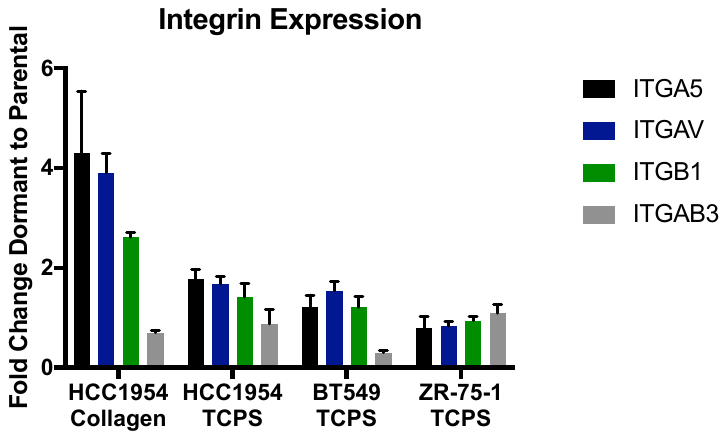

**Figure S7. Integrin expression of dormant cells determined by qPCR.** Each cell line was analyzed after 28 days of culture in serum-free medium on the substrate noted in the figure. Data is reported as fold change from the parental cell line.

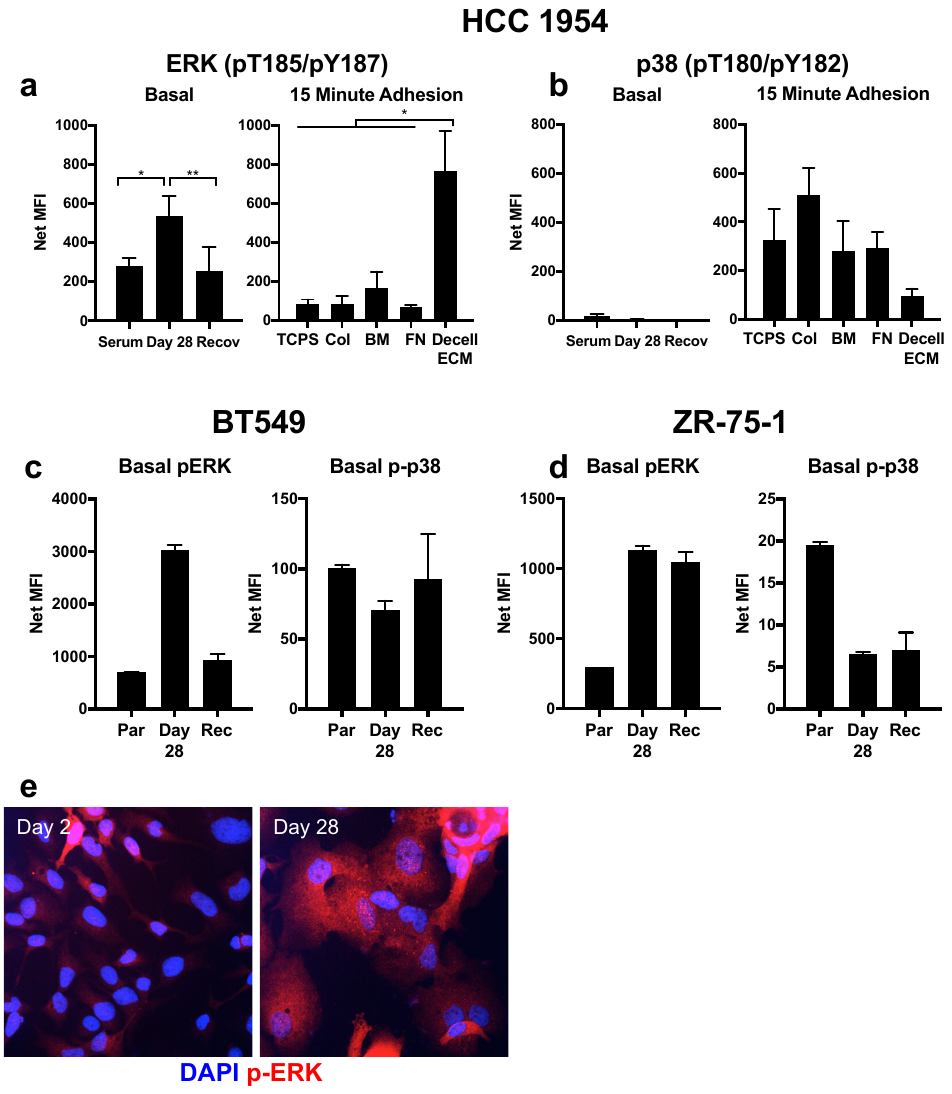

**Figure S8. ERK mediates survival in other cell lines.** a-b. Measurement of phospho-proteins via bead-based ELISA in HCC 1954 cells. Left, cells grown in serum and latent for 28 days on collagen and right, during adhesion to ECMs. Serum: cells in serum for 48h, Day 28: latent cells cultured for 28 days, Recov: latent cells recovered for 7 days. Col: collagen, BM: bone marrow, FN: fibronectin, Decell ECM: decellularized matrices obtained from latent HCC 1954 cells cultured on collagen coverslips for 28 days. c-d. Measurement of basal phospho ERK (pT185/pY187) and p38 (pT180/pY182) in parental, serum starved, and recovered BT549 (c) and ZR-75-1 (d) cells. Par: parental cells in serum, Day 28: dormant, Rec: recovered for 7 days. e. Immunofluorescence confirmation of elevated phospho-ERK (Thr202/Tyr204) in HCC 1954 serum starved on collagen at day 2 and 28. Scale: 100μm.

**Supplemental Tables**

Supplemental Table 1. Inhibitors and activators used in experiments

| **Target** | **Product** | **Company** | **Working Concentration** |
| --- | --- | --- | --- |
| E64 | 5208 | Tocris | 5 μM |
| ERK | FR180204 | Sigma-Aldrich | 20 μM |
| FAK | FAK inhibitor 14 | Sigma-Aldrich | 10 μM |
| Soluble Fibronectin | FC010 | Millipore | 10 μg/ml |
| Integrin α_5_ | AF1864 | R&D Systems | 1 μg/ml |
| Integrin α_v_ | Cilengitide | Apex Biotechnology | 100 μM |
| Integrin β_1_ | MAB17781 | R&D Systems | 1 μg/ml |
| MEK | PD0325901 | Sigma-Aldrich | 10 μM |
| MMP (pan) | GM6001 | Millipore | 25 μM |
| ROCK | Y-27632 | Millipore | 10, 1, 0.1 μM |
| TGFβ(pan) | AB-100-NA | R&D Systems | 20 μg/ml |
| TGFBR1 | LY-364947 | Sigma-Aldrich | 5 μM |

Supplemental Table 2. Antibodies used for immunofluorescence

| **Target** | **Company** | **Product** | **Dilution** |
| --- | --- | --- | --- |
| BrdU | Abcam | ab8039 | 2 μg/ml |
| Collagen I | Abcam | ab6308 | 1:200 |
| EdU | Life Technologies | Click-iT EdU | Per manufacturer’s protocol |
| ERK (phospho Thr202/Tyr204) | Cell Signaling | 4370 | 1:200 |
| Fibronectin | BD Biosciences | 563100 | 1:200 |
| Ki67 | Abcam | ab16667 | 1:200 |
| Laminin (pan) | Abcam | ab11575 | 1:100 |
| Osteopontin | Abcam | ab8448 | 1:1000 |
| p38 (phospho Thr180/Tyr182) | Cell Signaling | 4511 | 1:800 |
| p21 | Abcam | ab7903 | 1:100 |
| Vitronectin | Abcam | ab13413 | 1 μg/ml |

**Supplemental Table 3. Integrin and CDKN Primers**

| Gene | Accession number | Position | Forward sequence, 5’-3’ | Position | Reverse sequence, 5’-3’ |
| --- | --- | --- | --- | --- | --- |
| ITGA5 | NM_002205.3 | 384 | GCTTCAACTTAGACGCGGAG | 559 | ACAGAGGTAGACAGCACCAC |
| ITGAv | NM_001144999.2 | 916 | TTCTGTAGCTGCCACTGACA | 1103 | CAAACCGTGCAAAGACCTCA |
| ITGB1 | NM_002211.3 | 2054 | CATCTGCGAGTGTGGTGTCT | 2262 | GGGGTAATTTGTCCCGACTT |
| ITGB3 | NM_000212.2 | 104 | CAACATCTGTACCACGCGAG | 350 | ACTGACTTGAGTGACCTGGG |
| CDKN1A (p21) | NM_000389.4 | 1021 | ATGAAATTCACCCCCTTTCC | 1194 | CCCTAGGCTGTGCTCACTTC |

**Supplemental Movie 1.** Rare cells divide after four weeks of serum free culture. Movie is HCC 1954 on collagen at day 28, length is 12 hours.
